# SARS-CoV-2-induced dysregulation in ADAR editing patterns persists post viral clearance in individuals with mild COVID-19

**DOI:** 10.1101/2025.06.09.658648

**Authors:** Aiswarya Mukundan Nair, Helen Piontkivska

## Abstract

Severe acute respiratory syndrome coronavirus 2 (SARS-CoV-2) infections remain a public health concern worldwide. Viral antigen triggered innate immune response leads to induction of interferons (IFNs) and interferon stimulated genes (ISGs) including ADAR1 p150 isoform, which edits adenosine (A) residues within double stranded RNAs in both the virus and the host. Such RNA editing mediated by ADARs plays a crucial role in innate immune responses during viral infections through modulation of host-virus interactions. Additionally, ADAR editing acts post-transcriptionally, and serves as a mechanism of dynamic regulation of transcriptome and proteome diversity. While evidence points to changes in ADAR editing during infection, we do not know whether editing targets change over the course of the infection. Here, we explored temporal changes in ADAR expression and editing patterns, across three distinct stages of SARS-CoV-2 infection. Furthermore, we examined whether infection-triggered dysregulation in ADAR editing persists or returns to pre-infection states post-viral clearance. We addressed this question by analyzing publicly available whole blood RNA sequencing samples from forty-five, age-matched individuals. The individuals selected had no documented comorbidities, developed mild COVID-19, and were sampled across three distinct stages of SARS-CoV-2 infection: pre-, mid-, and post-infection. Our results demonstrate dynamic changes in ADAR expression and editing across the three stages. We further identified unique editing sites resulting from SARS-CoV-2 infection, across all three stages of infection, within genes involved in immune response pathways. Noteworthy, genes within neutrophil degranulation pathway appear to be edited, suggesting they may play a role in inflammation and sequelae observed post-SARS-CoV-2 infection. Our results demonstrate a consistent trend of elevated ADAR expression and reduced overall ADAR editing within each individual mid-infection. Subsequently, in some post-infection samples ADAR expression returns to approximately pre-infection levels, while in others it remains dysregulated. These differences may be contributing to heterogeneity in disease outcomes seen in individuals post-SARS-CoV-2 infection.

## Background

Severe Acute Respiratory Syndrome Coronavirus-2 (SARS-CoV-2), a previously unknown coronavirus, emerged in December 2019, and rapidly spread resulting in a pandemic (Abdelrahman et al., 2020; Holmes et al., 2021) and is associated with significant morbidity and mortality worldwide (*WHO COVID-19 Dashboard*, 2025). Coronaviruses cause respiratory, gastrointestinal, and neurological illnesses (Dessau et al., 2001; Parasa et al., 2020; V’kovski et al., 2021). While most coronaviruses commonly found in the human population are associated with mild illness, Middle East Respiratory Coronavirus (MERS-CoV), Severe Acute Respiratory Syndrome Coronavirus (SARS-CoV), and SARS-CoV-2 have been shown to cause diseases with a wider spectrum of severity (Corman et al., 2018; C. Huang et al., 2020). Clinical manifestations of Coronavirus disease 2019 (COVID-19), the disease caused by SARS-CoV-2 infection, can range from mild disease with symptoms such as sore throat, fever, dyspnoea, fatigue, myalgia and cough (Baj et al., 2020; Chen et al., 2020; C. Huang et al., 2020), to severe manifestation, when life-threatening conditions such as pneumonia and acute respiratory distress syndrome (ARDS) (Lamers & Haagmans, 2022) may develop. Additionally, a significant proportion of recovered individuals experience persistent symptoms or develop new symptoms months to years post-infection, referred to as the post-acute sequelae of COVID-19 (PASC) (C. Huang et al., 2020; Parotto et al., 2023; Wang et al., 2020).

Upon SARS-CoV-2 infection, the positive single stranded RNA (+ssRNA) is replicated within the host cell, forming double-stranded RNA (dsRNA) replication intermediaries. These dsRNAs activate innate immune sensors resulting in the production of interferons (IFNs) and other proinflammatory cytokines (Samuel, 2023). Interferon production subsequently results in transcription of interferon stimulated genes (ISGs) including an isoform of ADAR1, ADARp150 (Samuel, 2023; Song et al., 2022). This ADARp150 shuttles between the nucleus and the cytoplasm, acting on both viral and host transcripts, thereby altering ADAR editing patterns (Ringlander et al., 2022; Samuel, 2023; Song et al., 2022). Moreover, increased levels of the constitutively expressed ADAR1 isoform, ADARp110, and subsequent changes in ADAR editing patterns have also been reported during viral infections (Nachmani et al., 2014), including SARS-CoV-2 infection (Peng et al., 2022; Picardi et al., 2022). Because of competitive interactions between the three ADARs (Cruz et al., 2020), an increase in ADAR1 expression during viral infection could also impact the expression and activity of other ADARs (Cenci et al., 2008; Savva et al., 2012a; Tsivion-Visbord et al., 2020).

The dual role of ADAR editing, as a key component of innate immune response and as transcriptome regulator (Eisenberg & Levanon, 2018; Piontkivska et al., 2021), makes it a potential factor contributing to symptoms observed both during and post-viral infection (Jin et al., 2024; Piontkivska et al., 2019; Tsivion-Visbord et al., 2020; Wales-McGrath et al., 2023). ADARs edit adenosines (A) residues to inosines (I), within coding and non-coding regions of transcripts (Bass, 2002), which are then interpreted as A-to-G changes by the cellular machinery. For consistency, here we refer to A-to-G (and complementary T(U)-to-C) nucleotide changes as ADAR editing. Depending on the genomic region incorporating these edits, ADAR mediated editing can influence multiple aspects of gene expression. For instance, within coding regions ADAR editing sites can introduce non-synonymous substitutions potentially influencing protein structure and function. Editing sites within non-coding regions can impact alternative splicing events, microRNA and circular RNA biogenesis and function, and RNA stability (Kapoor et al., 2020; Kawahara et al., 2008; Nishikura, 2016; Slotkin & Nishikura, 2013).

ADAR mediated editing has been identified in several viral RNAs, including Zika virus (Piontkivska et al., 2017), Influenza virus (Suspène et al., 2011), Rift Valley fever virus (Suspène et al., 2008), hepatitis delta virus (Gélinas et al., 2011), human immunodeficiency virus type 1 (HIV-1) (Clerzius et al., 2009), and human respiratory syncytial virus (Martínez & Melero, 2002), where it could have a pro- or anti-viral effect depending on virus- and host-specific factors (Piontkivska et al., 2021; Samuel, 2011). In addition to editing of viral genome, viral infection-triggered changes in host ADAR editing patterns have been reported in multiple viral infections (Piontkivska et al., 2021), including during Zika virus infection (Piontkivska et al., 2019; Wales-McGrath et al., 2023), reovirus infection (Hood et al., 2014; Tariq & Piontkivska, 2024), and CMV infection (Wales-McGrath et al., 2023). Emerging evidence suggests altered ADAR editing patterns in the host during SARS-CoV-2 infection (Crooke et al., 2021b, 2021a; Merdler-Rabinowicz et al., 2023). Crooke et al. (2021a) identified reduced editing of Alu elements in individuals with severe COVID-19 and in dendritic cells infected with SARS-CoV-2. They proposed accumulation of unedited Alu dsRNAs in heightened inflammatory response associated with severe COVID-19 (Crooke et al., 2021a). In another study using normal human bronchial epithelial cells (NHBE) and biopsy samples, Crook et al. (2021b) replicated the previous results of reduced Alu editing and suggested under editing of Alu dsRNA as potent activators of IRF and NF-kB signaling-associated inflammatory response (Crooke et al., 2021b). On the other hand, Merdler-Rabinowicz et al. (2023), using nasopharyngeal swab and whole-blood samples of patients with COVID-19, identified a significant increase in global and site specific editing including in protein coding genes involved in immune regulation, and proposed such changes to have long-term physiological consequences (Merdler-Rabinowicz et al., 2023). In a recent study Jin et al. (2024) showed that altered ADAR editing in multiple ocular tissues in response to SARS-CoV-2 infection is associated with ophthalmic manifestations of COVID-19 (Jin et al., 2024).

While we already have some evidence that SARS-CoV-2 triggers changes in ADAR editing of host transcripts (Crooke et al., 2021b, 2021a; Jin et al., 2024; Merdler-Rabinowicz et al., 2023), the trajectory of temporal changes in ADAR expression and ADAR editing across distinct stages of SARS-CoV-2 infection within the same individuals is unknown. Additionally, whether SARS-CoV-2 triggered changes in ADAR editing patterns in the host persists or returns to pre-infection state during post-viral clearance is poorly understood. Here we explore temporal changes in ADAR editing patterns across pre-, mid-, and post-stages of SARS-CoV-2 infection, within the same individuals using whole-blood RNA sequencing samples obtained from forty-five age-matched, physically fit individuals without documented comorbidities who developed mild COVID-19 (BioProject PRJNA815324, COVID-19 Health Action Response for Marines, CHARM study) (Sauerwald et al., 2022).

## Results

The original dataset from the CHARM study (Sauerwald et al., 2022) has been used across multiple studies to investigate alterations in various molecular processes during SARS-CoV-2 infection (Z. Zhang et al., 2023). We extend the analysis from previous studies using a subset of the same dataset to identify dysregulation of ADAR editing patterns across three distinct stages of SARS-CoV-2 infection within the same individuals. Given that the original dataset included individuals that were often sampled only once, we selected a subset of forty-five individuals for whom samples were available across the three stages of infection (Supplementary Table 9), allowing us to investigate temporal changes in ADAR expression and editing patterns and to determine whether infection induced dysregulation persists or returns to pre-infection levels post-viral clearance.

### Expression dynamics of ADAR genes and isoforms

Pairwise differential gene expression analysis, between pre- and mid-infection samples and between pre- and post-infection samples was performed using DESeq2 package (Love et al., 2014) to identify the effects of SARS-CoV-2 infection on the host transcriptome. One thousand seventeen genes were found to be differentially expressed mid-infection compared to pre-infection (Supplementary Table 1A) according to filtering criteria of log_2_Fold Change > |0.58| (Fold Change > 1.5) and an adjusted p value < 0.05. Of these, 665 genes were upregulated, and 354 genes were downregulated. We further explored whether interferon stimulated genes (ISGs) including ADAR1 are differentially expressed mid-SARS-CoV-2 infection in our subset, by comparing the list of differentially expressed genes to a list of 329 ISGs from Schoggins et al. (Schoggins et al., 2011) that includes ADAR1. Out of the 1017 genes differentially expressed mid-infection (compared to pre-infection), 97 were ISGs (Supplementary Table 1C), however ADAR1, although upregulated mid-infection (log_2_Fold Change = 0.487, adjusted p value = 3.02E-06), did not meet the filtering criteria for differential expression.

Further, Reactome (Fabregat et al., 2017) was used to identify pathways in which the differentially expressed genes were overrepresented. Only pathways that met the filtering criteria of an FDR =< 0.05 and p value < 0.05 were considered overrepresented. Mid-infection differentially expressed genes (compared to pre-infection) were enriched in immune system-related pathways, including “Interferon alpha/beta signaling”, “Complement cascade”, “Signaling by the B Cell Receptor (BCR)”, and “Immunoregulatory interactions between a Lymphoid and a non-Lymphoid cell”, in addition to cell cycle-related pathways, including “Separation of Sister Chromatids” and “G2/M Checkpoints”, and other disease-related pathways. Full list of significantly overrepresented Reactome pathways among genes differentially expressed mid-infection are listed in the Supplementary Table 2A.

Differential gene expression analysis between pre- and post-infection samples identified five genes (Supplementary Table 1B), namely Granzyme B (GZMB), small nucleolar RNA – 118 (SNORD118), Ubiquinone oxidoreductase subunit A7 (NDUFA7), zinc finger and BTB domain containing 32 (ZBTB32), and small nucleolar RNA, H/ACA box 70 (SNORA70). Despite the low number of differentially expressed genes post-infection, Reactome pathway overrepresentation analysis identified these genes to be enriched (FDR < 0.05 and p value < 0.05) in signal transduction (“Signaling by NOTCH”) and disease associated pathways, including “Signaling by ALK in cancer”. However, using a less stringent cutoff of FDR =< 0.1 and p value < 0.05 identified enrichment of other pathways, including metabolism pathways (“Vitamin E”), disease pathway (“Nuclear events stimulated by ALK signaling in cancer”), and programmed cell death pathways (“Activation, myristolyation of BID and translocation to mitochondria”) (Supplementary Table 2B). Using a stricter filtering criteria of log_2_Fold Change > |1| (Fold Change > 2) and an adjusted p value < 0.05, showed 399 and 3 genes to be differentially expressed mid- and post-infection, respectively, compared to pre-infection with similar pathways as observed above (Supplementary Table 1D,E and Supplementary table 2C).

Using the relatively strict cut-offs described above, ADARs were not found to be differentially expressed among the ISGs, mid-SARS-CoV-2 infection in our subset of patients. Nonetheless, considering the nuanced and non-linear relationship between ADAR editing activity and its expression, where even a relatively minor change in expression may lead to functionally significant protein consequences downstream (Jacobs et al., 2009), we examined dysregulation in the expression of ADARs measured as Transcripts per million (TPM) normalized values. Comparisons were made between pre- and mid-infection samples, and between pre- and post-infection samples, to identify changes in the levels of ADARs mid-infection and to examine whether the changes observed mid-infection return to pre-infection levels post-viral clearance. Our analysis revealed a significant increase in ADAR1 expression in mid-infection compared to pre-infection (paired t-test: p value = 1.7 x 10^-6^). However, this increase in ADAR1 expression observed mid-infection was not retained post-infection (paired t-test: p value = 0.96), and no significant change was observed in the expression levels of ADAR1 post-infection when compared to pre-infection, suggesting that ADAR1 expression levels return to nearly pre-infection levels, post viral clearance (Fig. 1A) (Supplementary Table 3A). We also examined the changes in expression of ADAR2 and ADAR3, both mid- and post-infection compared to pre-infection. ADAR2 expression was elevated mid-infection compared to pre-infection, however, this change did not achieve statistical significance (paired t-test: p value = 0.07). Furthermore, there was a marginal decrease in ADAR2 expression post infection compared to pre-infection (Supplementary Table 3B), however, no significance was achieved (paired t-test: p value = 0.22) (Fig. 1B). ADAR3 was minimally expressed across all three stages of infection and no significant change was observed in its expression, both mid-(paired t-test: p value = 0.36) and post-infection (paired t-test: p value = 0.5) compared to pre-infection (Fig. 1C) (Supplementary Table 3C).

**Figure 1:**
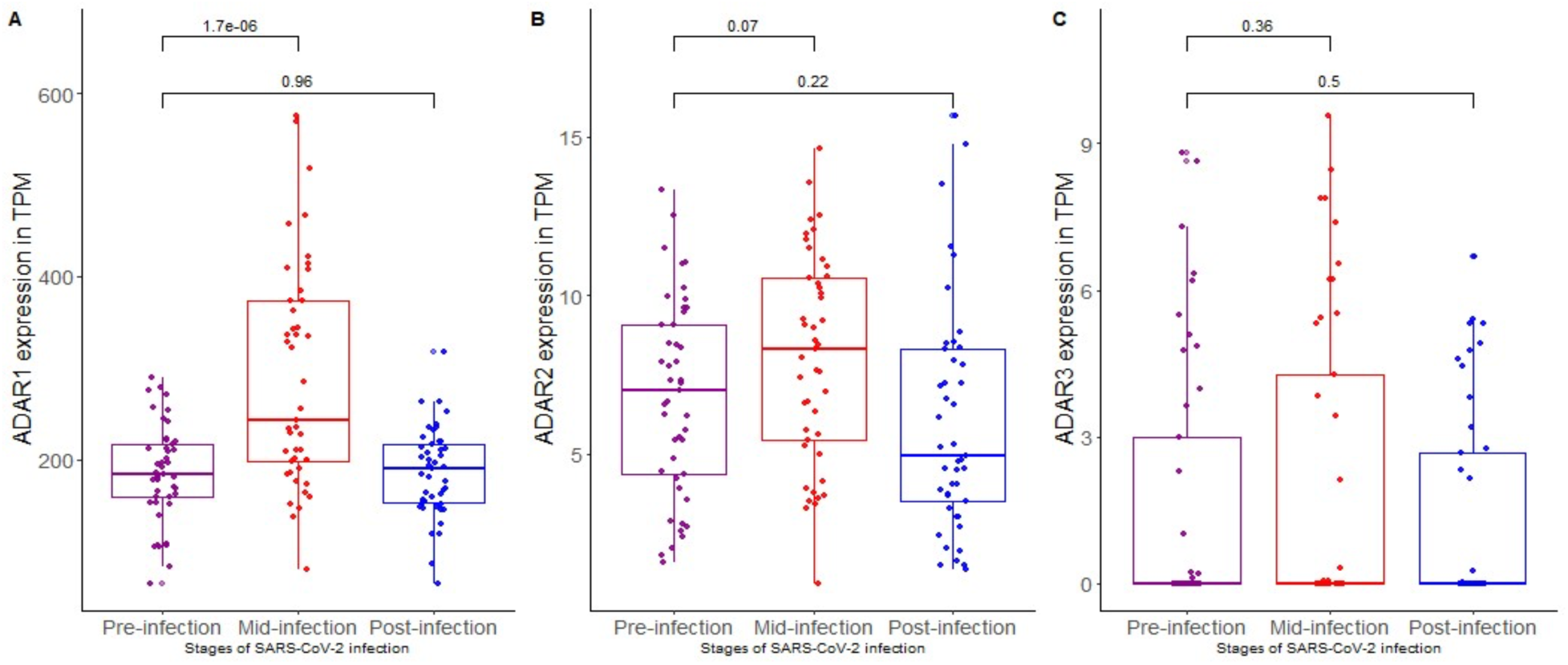
Expression dynamics of ADAR genes, in transcripts per million (TPM), across three distinct stages of SARS-CoV-2 infection: pre-, mid-, and post-infection. (A) ADAR1 (ADAR) expression increased significantly mid-infection compared to pre-infection (paired t-test, p value = 1.7 x 10^-6^). However, this increase in expression is not retained in the post-infection stage, and no significant difference in ADAR expression is seen between pre- and post-infection samples (paired t-test p value = 0.96). (B) ADAR2 (ADARb1) expression increased mid-infection (median TPM 8.309; paired t-test, p value = 0.07) and decreased minimally post-infection (median TPM 4.958; paired t-test, p value = 0.22) compared to pre-infection (median TPM 6.991), although the observed differences did not achieve statistical significance in both comparisons. (C) ADAR3 is minimally expressed across all stages of infection and showed no significant change mid-(paired t-test, p value = 0.36) or post-infection (paired t-test, p value = 0.5) compared to pre-infection

We further analyzed transcript level expression of ADAR1 using DESeq2 normalized counts, specifically looking for changes in expression of constitutively expressed ADAR1 isoform ADARp110 and interferon-inducible isoform ADARp150, both mid- and post-infection compared to pre-infection. Expression level of ADARp110 was significantly higher mid-infection compared to pre-infection (paired t-test using DESeq2 normalized counts: p value = 0.006, DESeq2 p-adj = 0.8906, log2FoldChange 0.6076). However, there were no significant changes in ADARp110 expression between pre- and post-infection samples, which returned to around pre-infection levels post viral clearance (paired t-test using DESeq2 normalized counts: p value = 0.98, DESeq2 p-adj = 1, log2FoldChange 0) (Fig. 2A). Expression level of ADARp150, although minimal, was increased mid-infection compared to pre-infection, however, this change did not reach statistical significance (paired t-test using DESeq2 normalized counts: p value = 0.31, DESeq2 p-adj 0.9643, log2foldChange 0.3360). Similar to ADARp110, post-infection levels of ADARp150 returned to nearly pre-infection levels (paired t-test using DESeq2 normalized counts: p value = 0.95, DESeq2 p-adj = 1, log2foldChange 0.0205) (Fig. 2A) (Supplementary Table 4A, B).

**Figure 2:**
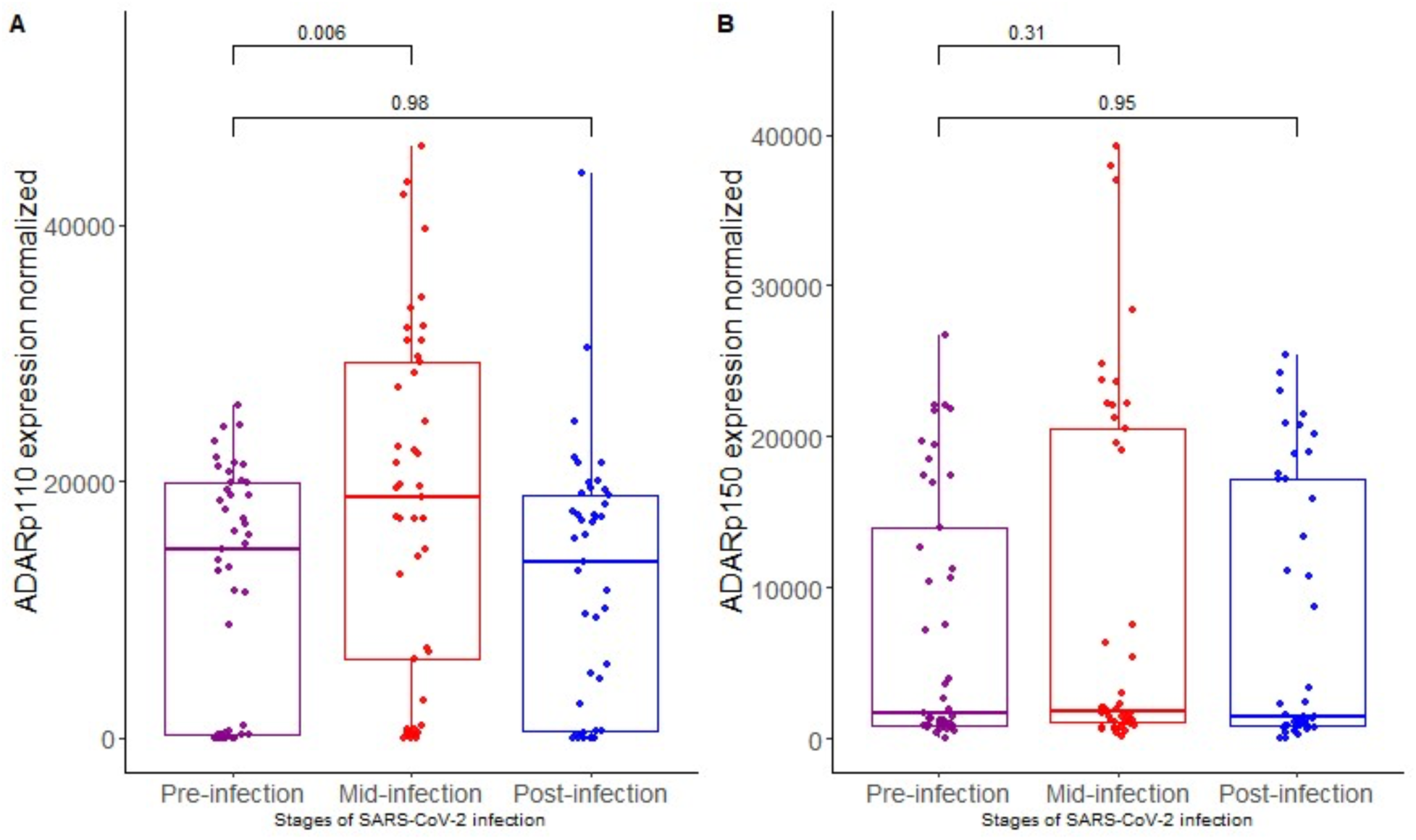
Changes in expression of ADAR isoforms, ADARp110 and ADARp150 as DESeq2 normalized counts, across mid-, and post-infection, compared to pre-infection. (A) ADARp110 expression is significantly increased mid infection compared to pre-infection (paired t-test, p value = 0.006), although this increase in expression is not retained post-infection, where no significant change is observed in the expression of ADARp110 when compared to pre-infection (paired t-test, p value = 0.98). (B) Transcript level analysis of ADARp150 showed that expression increased mid-infection compared to pre-infection, although statistical significance was not achieved (paired t-test, p value = 0.31). ADARp150 expression showed no change post-infection compared to pre-infection (paired t-test p value = 0.95).

### Alterations in number and genomic distribution of ADAR editing sites

To analyze the effects of elevated ADAR expression (both ADAR1 and ADAR2) on host ADAR editing patterns, we first examined changes in total number of putative ADAR edits that included A-to-G and T-to-C substitutions, excluding single nucleotide polymorphisms (SNPs) and potential polymorphic sites. We first identified all possible base substitutions from GATK generated VCF files including A-to-G, T-to-C, A-to-C, A-to-T, C-to-A, C-to-G, C-to-T, G-to-A, G-to-C, G-to-T, T-to-A, and T-to-G substitutions. Studies have identified A-to-G and T(U)-to-C substitutions, representing potential ADAR edited sites, as the most abundant type of substitution in metazoans (Nishikura, 2010a). Consistent with these findings, A-to-G and T-to-C substitutions were the most abundant form of substitutions across all three stages of infection. (Supplementary Figure 1A). Overall, the number putative ADAR edits increased mid-infection (average of 68151 ± 3042 edits per sample) compared to pre-infection (average of 62489 ± 3717 edits per sample), however, this increase was not statistically significant (paired t-test, p value = 0.15). Similarly, no significant change was observed in the total number of putative ADAR edits post-infection (average of 62193 ± 3187 edits per sample) compared to pre-infection (paired t-test, p value = 0.94). (Table 2) (Supplementary Figure 1B). Nonetheless, the total number of putative ADAR edits, normalized for per sample read depth, varied within individuals when comparing the total number of ADAR edits mid- and post-infection to their respective pre-infection numbers (Fig. 3A) (Supplementary Table 5).

**Figure 3:**
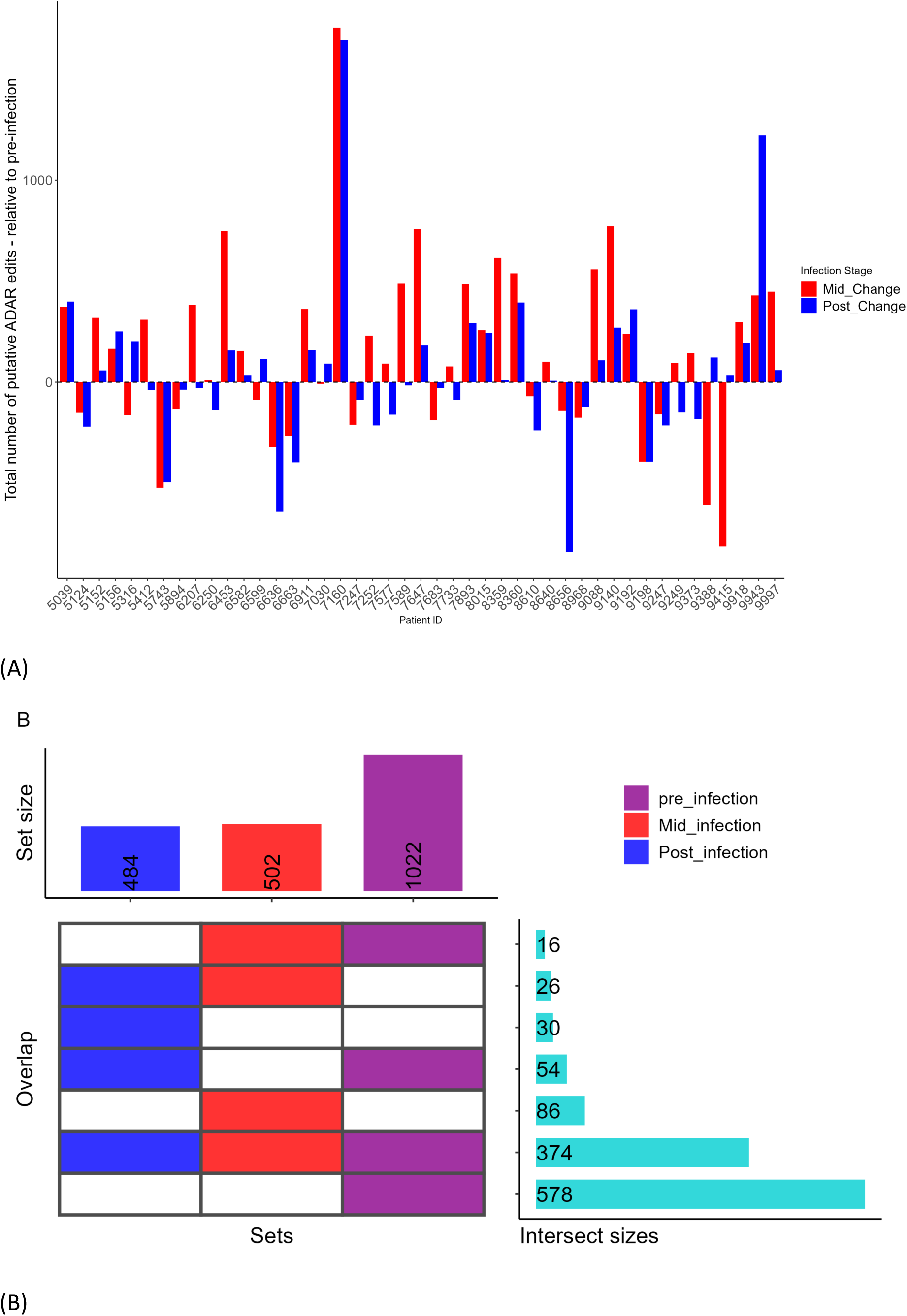
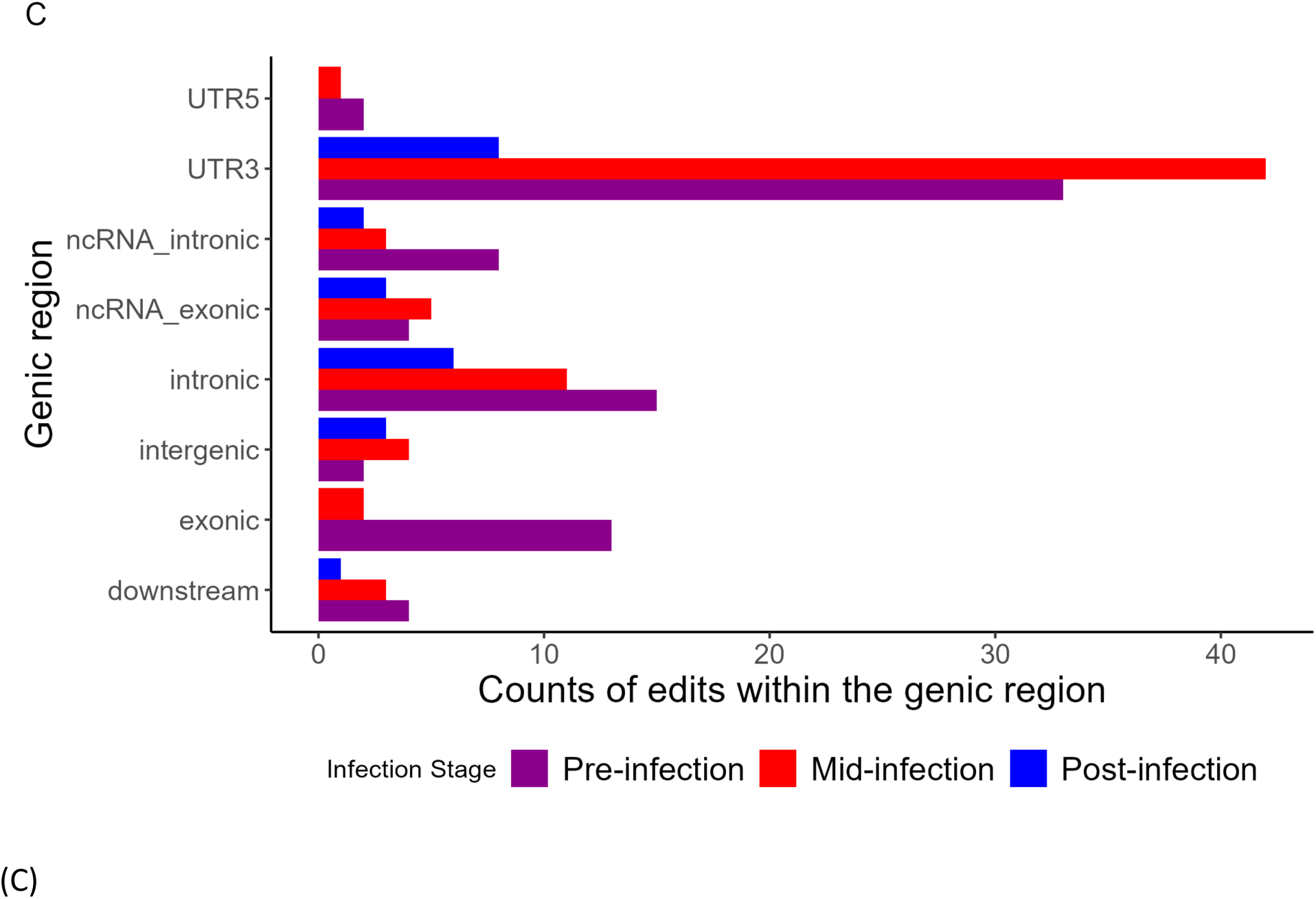
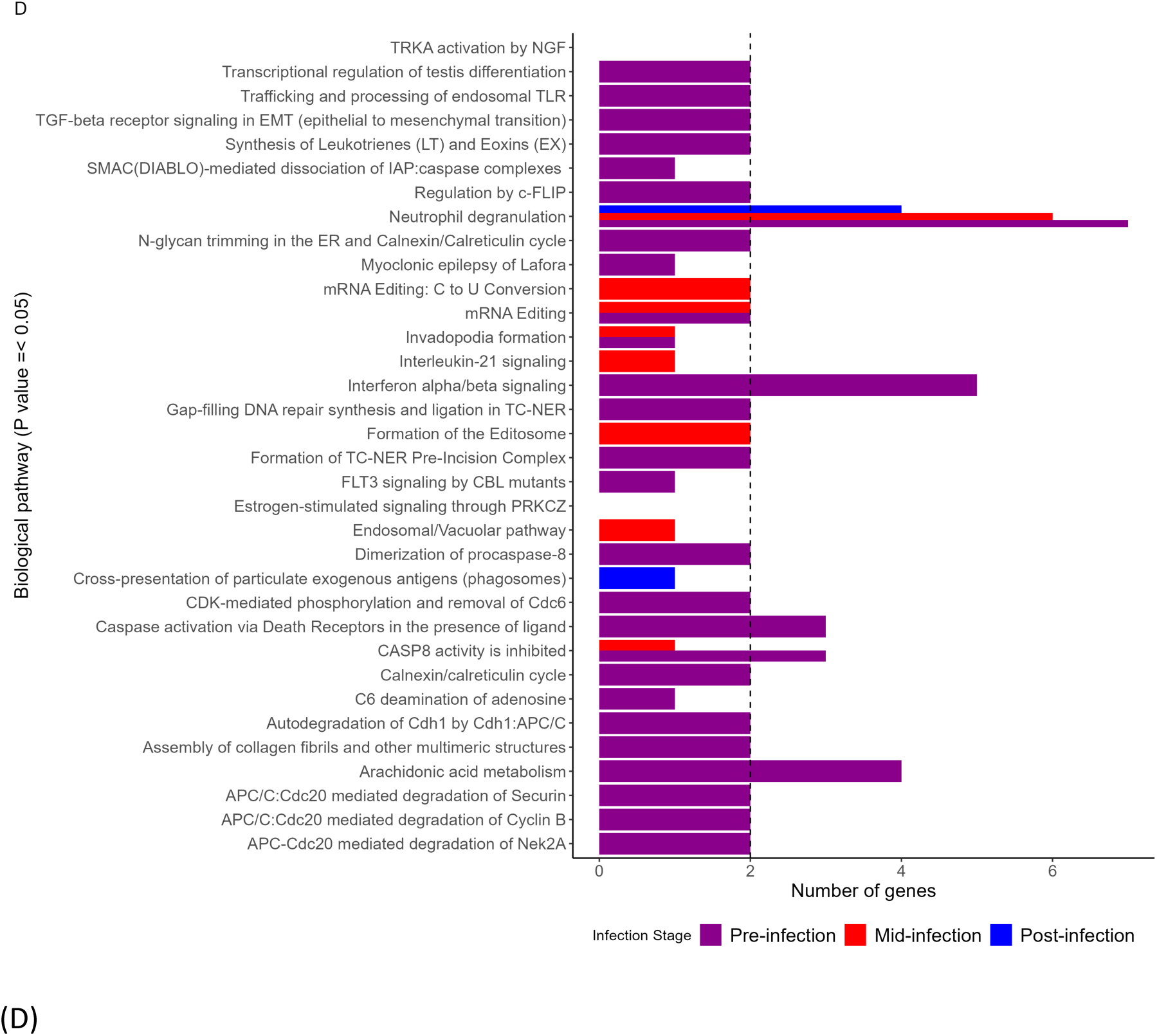
Changes in the number of ADAR editing sites including within individual changes in total number of putative ADAR edits, number of unique ADAR edits, distribution of unique ADAR edits across genic regions, and pathway enrichment analysis of genes incorporating unique edits. (A) Total number of putative ADAR edits varied within each individual mid- and post-infection relative to respective pre-infection values. (B) The upset plot shows the number of consistently and uniquely edited sites across pre-, mid-, and post-infection stages of infection. The plot also shows the number of sites shared between different infection stages. (C) Unique editing sites by genic regions and (D) Pathways overrepresented among genes incorporating unique editing sites pre-infection, mid-infection, and post-infection.

**Table 1:** K-means clustering analysis on overall ADAR editing levels pre- and post-infection clustering analysis using pre- and post-infection overall ADAR editing values resulted in two different groups of patients. Patients from the first group had persistent dysregulation in overall ADAR editing levels post-infection, while overall ADAR editing levels of patients from the second group returned to around pre-infection levels post viral clearance.

| K-means group | Number of individuals within each group | Average pre-infection overall ADAR editing levels | Average post-infection overall ADAR editing levels |
| --- | --- | --- | --- |
| **Group 1**<br>**(Dysregulated editing)** | 12 | 0.6222 | 0.4290 |
| **Group 2**<br>**(Minimal difference in editing)** | 33 | 0.2514 | 0.2561 |

**Table 2:** Characteristics of a subset of individuals from BioProject PRJNA815324 (CHARM dataset) used in this study. A total of 45 individuals were selected with whole-blood RNA sequencing data available at three distinct stages of SARS-CoV-2 infection. The patients were age-matched (19.1 ± 1.9 years), physically fit, mostly Caucasians with a male to female ratio of 4:1. Values for the number of ADAR editing sites and overall ADAR editing levels are represented as mean and standard error of mean (SEM).

| SARS-CoV-2 infection stage | Total number of samples (N=135) | Days post T00 | Severity of COVID-19 | Average number of aligned reads per sample | Mean number of ADAR editing sites (A-to-T and T-to-C) per sample for each stage of infection (+/-) SEM | Mean overall ADAR editing levels per sample for each stage of infection (+/-) SEM |
| --- | --- | --- | --- | --- | --- | --- |
| Pre-infection | 45 | T00 | NA | 570,67,797 | 62489 ± 3717 | 0341603 ± 0.02541721 |
| Mid-Infection | 45 | T36 | Mild | 521,61,062 | 68151 ± 3042 | 0.2817583 ± 0.01892626 |
| Post-infection | 45 | T56-T105 | NA | 515,56,269 | 62193 ± 3187 | 0.2994999 ± 0.02119264 |

Next, we analyzed whether SARS-CoV-2 infection introduces unique ADAR editing sites mid- and post-infection (compared to pre-infection), or results in loss of existing (sites present pre-infection) ADAR editing sites, by identifying sites uniquely edited pre-, mid-, and post-infection. Towards this goal we first identified sites consistently edited across each distinct stage of infection. Consistently edited sites included those A-to-G and T-to-C substitutions, with a GATK mapping quality of 40, had 100 or more aligned reads, with an editing level of 2% or more, and were present in at least 20% of samples for any given stage of infection. While 1022 sites were identified as consistently edited pre-infection, the number of consistently edited sites decreased mid- and post-infection with 502 and 484 sites respectively (Fig. 3B) (Supplementary Table 6A). Consistently edited sites were further compared to identify the editing sites unique to the three stages of infection. 578 sites were found to be uniquely edited in pre-infection samples, while 86 and 30 sites were uniquely edited in mid- and post-infection samples, respectively (Supplementary Table 6B). Unique ADAR editing sites were further annotated using REDIportal database to identify previously confirmed editing sites (within the list of unique edits) and to identify genes and genomic regions incorporating these unique edits. Out of 578 ADAR editing sites unique to pre-infection samples, 81 (Supplementary Table 6C) were confirmed editing sites present in the REDIportal. Similarly, out of 87 sites unique to mid-infection, 71 (Supplementary Table 6D) were confirmed editing sites present in the REDIportal. For sites unique to post-infection, 23 (Supplementary Table 6D) out of 30 were confirmed editing sites present in the REDIportal.

We further identified genic regions mapped to confirmed unique editing sites (present in REDIportal) across the three stages of infection. Pre-infection unique sites were mostly in the 3’UTR (∼41%), intronic (∼19%), and exonic (∼16%) regions, while the rest were in 5’UTR, ncRNA exonic, ncRNA intronic, intergenic, and downstream regions. Similarly, mid-infection unique sites were mostly in the 3’UTR (∼60%), intronic (∼15%), and exonic regions (∼3%), and the rest were in ncRNA exonic, ncRNA intronic, intergenic, 5’UTR, and downstream regions. While there were no unique sites in the exonic and 5’UTR regions post infection, the highest number of unique sites were in 3’UTR (∼35%) regions, followed by intronic (∼26), noncoding exonic, and intergenic regions (∼13%), while the rest were in the ncRNA intronic, intergenic and downstream regions (Fig 3C).

Subsequently, pathway overrepresentation analysis using Reactome of genes with confirmed unique editing sites identified pathway unique to a stage of infection as well as those present across multiple infection stages. The immune system pathway, specifically, “Neutrophil degranulation”, was present across all three stages of infection, while RNA metabolism (“mRNA editing”), programmed cell death (“CASP8 activity is inhibited”), and extracellular matrix organization pathways (“Invadopodia formation”) were overrepresented pre- and mid-infection. Notably, multiple pathways were uniquely overrepresented across distinct stages. For instance, “Interferon alpha/beta signaling”, “Caspase activation via death receptor in the presence of a ligand”, and “Arachidonic acid metabolism” were overrepresented pre-infection. On the other hand, “mRNA editing: C to U conversion”, “Formation of Editosome”, and “Interleukin-21 signaling” were overrepresent mid-infection, while “Cross-presentation of particulate exogenous antigens (phagosomes)” and “Estrogen-stimulated signaling through PRKCZ” were uniquely overrepresented post-infection (Fig 3D). Full list of significantly overrepresented pathways among genes with confirmed unique editing sites are listed in Supplementary Table 7.

### Changes in overall ADAR editing levels

To further investigate global changes in ADAR editing levels mid- and post-SARS-CoV-2 infection compared to pre-infection, we computed overall ADAR editing levels for each sample. Overall ADAR editing levels were defined as the ratio of all the edited reads from REDIportal identified ADAR editing sites in a sample to the total the number of reads aligning to these sites (Breen et al., 2019; Tan et al., 2017). First, we examined cumulative changes in overall ADAR editing levels mid- and post-infection compared to pre-infection, aiming to understand changes within the entire patient cohort. Our analysis showed a statistically significant decrease in overall ADAR editing level mid-infection compared to pre-infection (paired t-test, p value = 0.0096). While overall ADAR editing levels remained lower post-infection compared to pre-infection, this change was not statistically significant (paired t-test, p value = 0.14) (Fig. 4A). Furthermore, we explored individual specific (within-patient) differences, to determine if the above observed changes in overall ADAR editing levels are replicated within each patient. Consistent with the above observation, overall ADAR editing levels were decreased in majority of individuals, mid-infection, represented by red bars in figure 4B, where each bar corresponds to a patient. Notably, few individuals retained the dysregulated levels of overall ADAR editing post-infection compared to pre-infection, while in others overall ADAR editing returned to around pre-infection levels (Fig. 4B), represented by blue bars in figure 4B.

**Figure 4:**
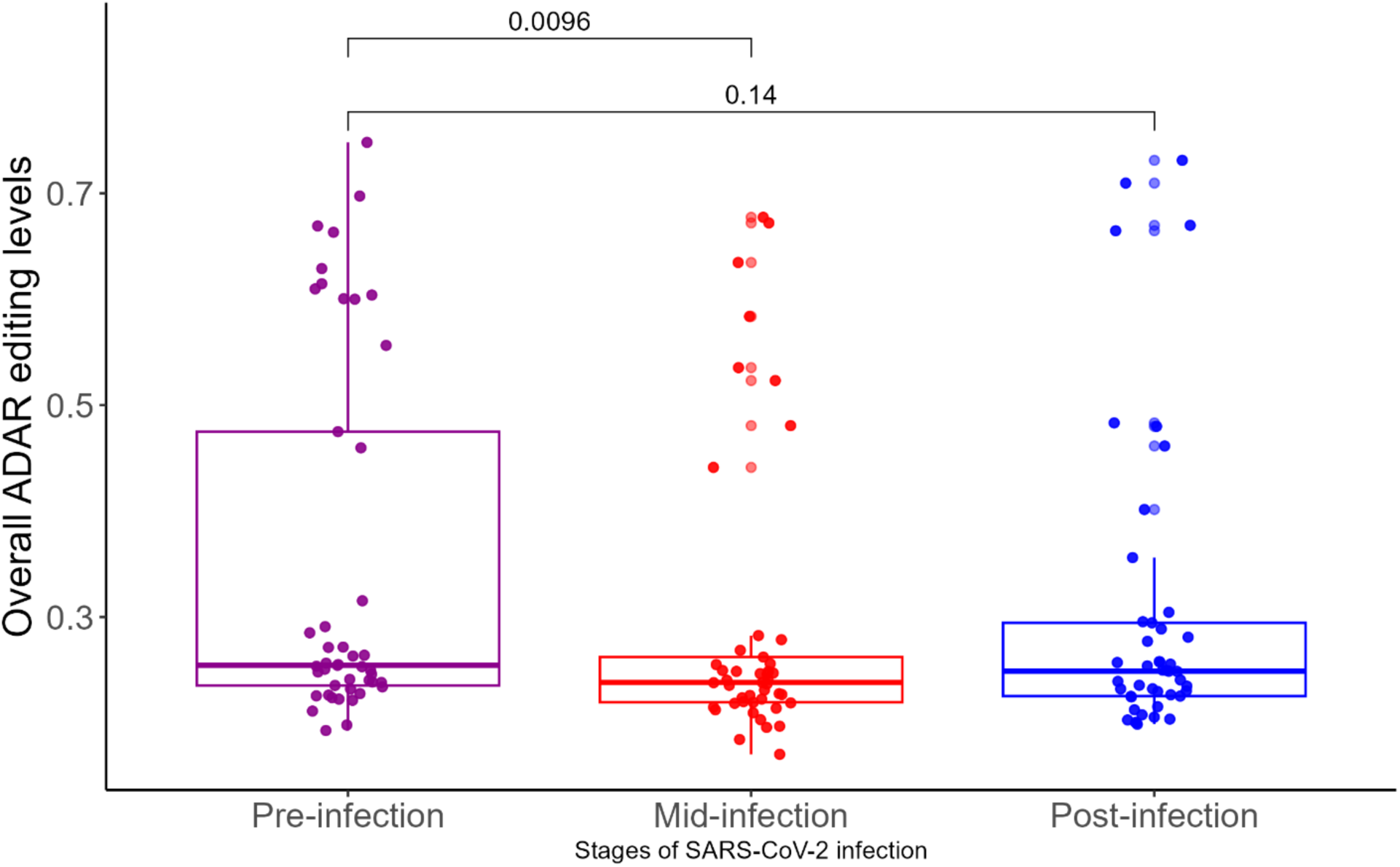
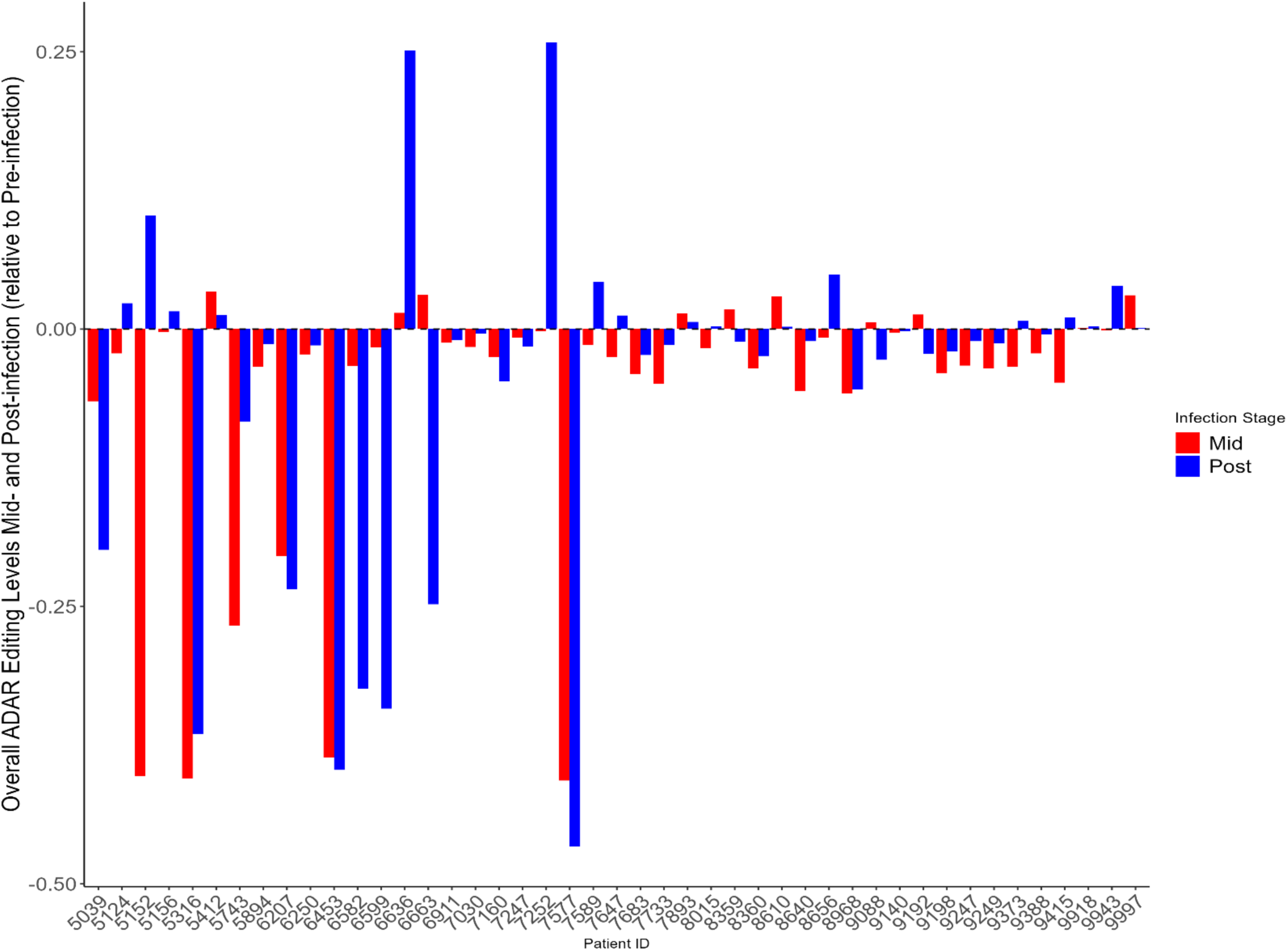
Alterations overall ADAR editing levels mid- and post-infection compared to pre-infection. Panel A and B show alterations in overall ADAR editing levels at cohort and individual specific levels respectively. Each bar in panel B represents an individual and values for mid-infection (red bars) and post-infection (blue bars) are represented relative to pre-infection overall ADAR editing values. (A) Overall ADAR editing levels showed statistically significant decrease mid-infection compared to pre-infection stage (paired t-test, p-value 0.011) and although overall ADAR editing levels remained decreased post -infection compared to pre-infection, the difference was not statistically significant (paired t-test, p value = 0.14). (B) Decreased levels of overall ADAR editing mid-infection compared to pre-infection was generally consistent across each patient. However, individual specific differences in overall editing levels are observed post-infection, where some individuals retained the dysregulated levels of overall ADAR editing while in others overall ADAR editing levels returned to around pre-infection levels post-infection.

Furthermore, to confirm the above observed patterns of overall ADAR editing post-infection – where some individuals retained the altered overall ADAR editing levels, while in others overall ADAR editing levels returned to around pre-infection levels – we performed K-means clustering analysis using pre- and post-infection overall ADAR editing level values within each individual. In line with the above observation, two clear patterns were observed, based on which individuals were divided into two groups. The first group consisted of 12 individuals, representing those with persistently dysregulated overall ADAR editing levels post-infection compared to pre-infection, with an average overall ADAR editing of 0.6222 and 0.4290, pre- and post-infection respectively. The second group consisted of 33 individuals, representing those with minimal difference in overall ADAR editing, with an average overall ADAR editing of 0.2514 and 0.2561 pre- and post-infection respectively (Table1). Supplementary Table 8 provides patient wise overall ADAR editing levels for both the groups. It is worth noting that the elbow method defined up to 4 potential groups of individuals in our dataset (Supplementary Figure 2), where group 1 represented individuals with lower pre-infection overall editing (0.3094) and higher post infection overall ADAR editing (0.4748), group 2 represented individuals with minimal differences in overall editing pre-(0.2461) and post-infection (0.2421). Group 3 represented individuals with a higher pre-infection overall editing (0.5892) and lower post-infection overall ADAR editing (0.2672), and group 4 individuals had higher baseline overall ADAR editing levels (0.6843) compared to the other groups and showed slight increase (0.6937) post infection (Supplementary Table 1). However, to align with our aim of identifying individuals with either dysregulated and/or minimal changes in overall editing levels post-SARS-CoV-2 infection, we defined 2 groups in K-means clustering analysis. We also performed K-means clustering using pre- and mid-infection overall ADAR editing values within each individual, using the elbow method, with up to two potential groups (Supplementary Figure 2A). However, this grouping was due to baseline (pre-infection) differences in overall ADAR editing levels and, in both groups, overall ADAR editing levels decreased mid-infection (Supplementary Table 2).

## Discussion

Viral infection-triggered induction of IFNs and subsequent changes in the expression and activity of the RNA editing enzyme ADAR play a significant role in shaping the outcomes for both the virus and the host through editing of viral and host transcripts (Di Giorgio et al., 2020; Nishikura, 2010). While several studies have identified the impact of host ADARs on SARS-CoV-2 genome (Di Giorgio et al., 2020; Peng et al., 2022; Picardi et al., 2022; Ringlander et al., 2022; Song et al., 2022), few studies have investigated the impact of SARS-CoV-2 infection triggered alterations in ADAR expression and activity on the host transcriptome (Crooke et al., 2021b, 2021a; Jin et al., 2024; Merdler-Rabinowicz et al., 2023). To the best of our knowledge, no studies have examined the temporal changes in ADAR expression and editing patterns pre-, mid-, and post-SARS-CoV-2 infection. Considering the dynamic and nuanced nature of ADAR editing (that can vary among individuals and stages of infection) (Jepson et al., 2011; Savva et al., 2012b), and the potential of dysregulated ADAR editing patterns to contribute to post-infection sequelae (Piontkivska et al., 2019; Tsivion-Visbord et al., 2020), we investigated temporal changes in ADAR expression and editing patterns at the pre-, mid-, and post-infection stages of SARS-CoV-2 infection, within the same individuals with comparable demographics (age-matched, without documented comorbidities). We further analyzed whether the dysregulation in ADAR expression and editing patterns, observed mid-infection, persists or returns to pre-infection state post-viral clearance. Our results demonstrate dynamic changes in ADAR expression and editing across three distinct stages of SARS-CoV-2 infection. We further provide evidence for unique changes in ADAR editing patterns resulting from SARS-CoV-2 infection, through either loss or introduction of editing sites unique to a stage of infection. Notably, our results show that ADAR expression and overall ADAR editing levels (representing global changes in editing) follow a similar trend among individuals, mid-infection. Specifically, there is a significantly increased ADAR expression and decreased overall editing mid-infection. On the other hand, despite ADAR expression returning to their pre-infection levels in post-infection samples, only some samples showed that overall ADAR editing levels returned to their pre-infection levels, while in others it remained dysregulated. This observation underscores how the nuanced nature of ADAR editing can vary even among individuals with comparable demographics. Finally, we propose persistent dysregulation in ADAR editing of host transcripts, including in genes involved in the neutrophil degranulation pathway, as one potential contributor to some of the symptoms observed in post-SARS-CoV-2 infection sequelae in a subset of individuals.

Transcriptomic analysis between pre- and mid-infection samples identified more than one thousand differentially expressed genes involved in immune system, cell cycle, DNA replication, cellular responses to stimuli, programmed cell death, vesicle-mediated transport, metabolism of proteins, transport of small molecules, hemostasis, and gene expression pathways, highlighting diverse cellular processes influenced by SARS-CoV-2 infection. Notably, the number of differentially expressed genes decreased substantially post-viral clearance. Five genes, GZMB, SNORD118, NDUFA7, ZBTB32, and SNORA70 met our criteria for differential expression. GZMB (granzymes), are cytolytic proteins released by cytotoxic T lymphocytes (CD8+ T cells) and natural killer cells during viral infection to eliminate infected cells. Persistent dysregulation of GZMB post-viral clearance is consistent with previous studies showing elevated levels of GZMB in individuals with PASC and in individuals recovering from COVID-19 (de Sousa Palmeira et al., 2023; Lin et al., 2022; Santa Cruz et al., 2023). Similarly, SNORD118 are small nucleolar RNAs primarily involved in ribosomal RNA modifications, and their expression levels are seen to be altered in lung diseases (Liu et al., 2023) as well as after influenza A viral infection (Zhuravlev et al., 2022), although their role if any during SARS-CoV-2 associated lung abnormalities is not yet understood. Interestingly, NDUFA7, (ubiquinone oxidoreductase) is an age-associated mitochondrial gene, whose expression decreases with age. NDUFA7 is seen to be repressed by SARS-CoV-2 infection and reflects common changes in aging and SARS-CoV-2 infection (Chow et al., 2021).

Although none of the three ADARs met our criteria for differential expression (log_2_Fold Change > |0.58|and an adjusted p value < 0.05), the expression of ADAR1, ADAR2 (measured as TPM normalized values) and ADAR1 isoforms (ADARp110 and ADARp150, measured as DESeq2 normalized values) increased mid-infection and returned to around pre-infection levels post-viral clearance. Notably, we found a significantly increased expression of constitutively expressed ADAR1 isoform ADARp110, while the expression of interferon inducible ADARp150 was only marginally increased. This was noteworthy, considering that ADARp150 expression is seen to be elevated during several viral infections and conditions of increased IFN in the cells (George & Samuel, 1999; Lamers et al., 2019; Salvetat et al., 2019), due to the presence of interferon stimulated response element (ISRE) (George et al., 2011; George & Samuel, 1999). However, the exact mechanism that leads to increase in ADARp110 expression during viral infection requires further studies (Nachmani et al., 2014; K. Zhang et al., 2022). Nonetheless, increased expression of ADARp110 in our dataset is consistent with a previous study by Picardi et al. (2022) that found increased ADAR editing in response to SARS-CoV-2 infection in Vero cells that lack the ability to produce IFNs and hence ADARp150, and suggested that the increase in ADAR editing could be due to the activity of ADARp110 (Picardi et al., 2022). Similar results have been reported by Peng et al., which suggested the role of host ADARp110 in addition to ADARp150 in editing of SARS-CoV-2 genome during infection (Peng et al., 2022). It is interesting to note that increased expression of ADARp110 was also observed by Nachmani et al. (2014) following human cytomegalovirus (HCMV) infection, who identified that this increase in ADARp110 expression was due to the activity of promoter1B, one out of three promoters of ADAR1 (1B, 1C and 2) that drive the expression of ADARp110 (Nachmani et al., 2014). Similarly, increased expression of ADARp110 has also been observed during reovirus infection (Tariq & Piontkivska, 2024) and enterovirus-D68 infection (K. Zhang et al., 2022). Findings from these studies, along with our results, highlight the importance of both ADARp110 and ADARp150 in editing of viral and endogenous host transcripts during viral infections. However, the exact mechanism that results in increased ADARp110 expression and whether it is specific to certain viruses requires further studies.

As mentioned above, expression of ADARp150 was only minimally increased in our samples mid-infection. A potential explanation for this could be the timing of sample collection. IFNs and ISGs (that includes ADArp150) exhibit dynamic expression throughout the course of infection, with peak expression often observed within first few hours after viral exposure in animal models, followed by a decline to undetectable levels (Chiale et al., 2022). Given that the precise time point of viral exposure is unknown for our dataset, it is possible that the levels of ADARp150 were higher earlier, but were decreasing by the time of sample collection. Furthermore, studies suggest that peripheral blood samples could not accurately represent IFN responses in other tissues (Chiale et al., 2022). Furthermore, ADAR2 levels increased minimally mid-infection, while there was no change in the expression of ADAR3, potentially due to its limited expression in tissue outside the nervous system (Melcher et al., 1996; Raghava Kurup et al., 2022).

While there was a marginal increase in the total number of putative ADAR editing sites during infection (without statistical significance), that number returned to around pre-infection levels post-viral clearance. Within individual comparisons of the total number of putative ADAR editing sites mid- and post-infection to the respective pre-infection numbers, edited sites differed within each individual, suggesting that increased expression of ADARs is translated to downstream difference in the number of ADAR edits, but to varying degrees (Fig. 3A). Nevertheless, we found a substantial decrease in the total number of consistently edited sites, representing those high-confidence sites that are shared across at least 20% of samples for a given stage of infection (refer to Methods for definition and filtering criteria of consistently edited sites), mid- and post-infection compared to pre-infection. Interestingly, the comparison of consistently edited sites across stages of infection identified editing sites that were unique to stages of infection, with the highest number of unique sites present in pre-infection followed by mid- and post-infection, demonstrating the nuanced and unique changes to ADAR editing landscape introduced by SARS-CoV-2 infection.

An appreciable number of pre-infection uniquely edited genes were in the exonic regions, with some genes, such as Interferon Induced Transmembrane Protein 2 (IFITM2), Post-GPI Attachment To Proteins 6 (PGAP6), Major Histocompatibility Complex, Class II, DQ Beta 1 (HLA-DQB1), and Ubiquitin C (UBC) incorporating multiple edits. A particularly notable among them is the transmembrane protein IFITM2, which is induced in response to interferons and functions to inhibit the fusion of viral and cellular membranes during viral infections (Diamond & Farzan, 2013; X. Zhao et al., 2019). IFITM2 has been identified to restrict the entry of a wide array of enveloped viruses including SARS-CoV-2 (I.-C. Huang et al., 2011; Prelli Bozzo et al., 2021). Two different non-synonymous substitutions (Chr11:308290 V/A and Chr11:308314 M/T) were found on the IFTM2 gene in our pre-infection samples; interestingly, non-synonymous substitutions on IFTM2 gene have also been previously reported during Zika infection (Wales-McGrath et al., 2023). While in the previous study, IFTM2 was found to be overedited during Zika infection, in our study this site was uniquely present in pre-infection samples suggesting potential loss of this site mid- and post-SARS-CoV-2 infection. Nonetheless, these previous reports, along with our results, suggest a potentially important role of non-synonymous ADAR editing on IFTM2 gene during viral infections. However, further studies are required to identify the impact of non-synonymous ADAR editing of IFTM2 gene on protein products (Wales-McGrath et al., 2023).

Although no unique exonic edits were present in post-infection samples, three genes incorporated exonic edits in mid-infection samples, namely, Adhesion G Protein-Coupled Receptor G3 (ADGRG3), Immunoglobulin Lambda Variable 1-51 (IGLV1-51), and bladder cancer associated protein (BLCAP) gene. Among these, BLCAP is a conserved tumor suppressor gene that is highly expressed in brain and B lymphocytes. It is known to undergo hyper-editing in various cancers and is associated with cell proliferation, cell cycle, and apoptosis (Paz et al., 2007; M. Zhao et al., 2016). In our dataset, we found nonsynonymous Q/R substitution at Chr22: 36147563 of the BLCAP gene. BLCAP is also seen to be edited during Zika infections (Wales-McGrath et al., 2023), and non-synonymous editing of BLCAP gene in Zika infection and during SARS-CoV-2 infection in our dataset could reflect potential alterations in cellular processes relevant to both cancer and viral infection; however, further investigations are warranted to draw concrete conclusions.

A majority of uniquely edited sites, across all stages of infection, were in the 3’UTR and intronic regions of the genes. An abundance of unique edits in these regions is not surprising considering the vast majority of editing occurs in the non-coding UTRs and introns (Gu et al., 2012; Hundley & Bass, 2010; Savva et al., 2012a). These non-coding regions regulate splicing, RNA stability, localization and translation through their interaction with RNA binding proteins and miRNAs, therefore, editing within these regions can have significant impact on gene expression (Aspden et al., 2023; Hong & Jeong, 2023). Importantly, dysregulation in these non-coding regions has been implicated in diseases including cancers (Hong & Jeong, 2023).

SARS-CoV-2 infection introduces significant alterations in innate and adaptive immune responses (Gelmez et al., 2022; Hajihasani et al., 2023), and a growing number of studies have identified that these alterations persist even after viral clearance to post-infection period (Ryan et al., 2022; Woodruff et al., 2023). In line with these findings, uniquely edited sites in our datasets, across all stages of infection (pre-, mid-, and post-infection), were enriched in innate and adaptive immune response pathways. Of particular importance is the neutrophil degranulation pathway, enriched among uniquely edited sites across all stages of infection. Degranulation is an important process mediated by neutrophils to eliminate intracellular and extracellular pathogens (Eichelberger & Goldman, 2020), associated with inflammation, it is seen in inflammatory disorders including asthma, acute lung injury, rheumatoid arthritis, and septic shock (Lacy, 2006). Transcriptomic and proteomic studies have identified significant activation of neutrophil degranulation pathway during SARS-CoV-2 infection and in individuals with PASC (Aschenbrenner et al., 2021; Long et al., 2022; Overmyer et al., 2021; Ryan et al., 2022), which indicates persistent inflammatory responses and immune dysregulation post-SARS-CoV-2 infection (Ryan et al., 2022). Noteworthily, neutrophil degranulation pathway was also enriched among mid-infection differentially expressed genes in our dataset which is consistent with previous studies. This suggests dysregulation of neutrophil degranulation associated genes at transcriptional and post-transcriptional levels potentially contributes to inflammation and symptoms observed both during and post-SARS-CoV-2 infection.

Notably, despite increased expression of ADAR1, we found a significant decrease in overall ADAR editing levels (a measure of global changes in ADAR editing within each sample) at mid-SARS-CoV-2 infection (compared to pre-infection). This decrease in overall editing was consistent across most of the individuals in our dataset. Given that the relationship between ADAR expression and ADAR editing is not always linear, and that increased expression of ADAR does not always lead to increased editing levels (Deffit & Hundley, 2016; Jacobs et al., 2009; Sapiro et al., 2020), this result is, perhaps, unsurprising. One possible explanation for this could be that increased ADAR1 interferes with the activity of other ADARs, as observed in previous studies (Orlandi et al., 2012; Cruz et al., 2020). For instance, a previous study in pediatric astrocytoma found decreased ADAR editing despite an increase in the expression of ADAR1 (specifically ADARp110), due to sequestration of ADAR2 by ADAR1 through heterodimer formation (Cenci et al., 2008). In a separate study, these authors identified decreased editing at Q/R site of the GluRB mRNA in glioblastoma cell line, despite overexpression of ADAR1, resulting from heterodimerization of ADAR1 and ADAR2. Moreover, regulation of RNA editing levels by various RNA binding proteins (RBPs) has been identified in Drosophila and various mammals. For instance, Sapiro et al. (2020) identified altered editing levels at multiple ADAR editing sites (site-specific regulation) following the knockdown of RBP Zn72D, which is distinct from global reduction in editing, which is seen in ADAR knockouts (Sapiro et al., 2020). The same study found that the loss of ZN72D resulted in significant decrease in ADAR protein levels, but not ADAR mRNA levels. These studies provide potential explanations for the decreased editing observed in our dataset. However, further studies are required to validate these findings and rule out the possibility of other presently unknown factors. Our comparison of overall ADAR editing levels between pre- and post-infection samples showed decreased editing, even though ADAR expression returned to around pre-infection levels; however, this change did not achieve statistical significance. Increasingly, studies have identified heterogeneity in various molecular processes among individuals post-SARS-CoV-2 infection (Le Bert et al., 2021; Ryan et al., 2022). Our analysis reveals persistent dysregulation in overall ADAR editing levels among a subset of individuals post-SARS-CoV-2 infection, while in others overall ADAR editing levels return around pre-infection levels post-viral clearance. Our results suggest that SARS-CoV-2 infection can lead to persistent changes in ADAR editing patterns even after viral clearance.

Our study focused on a relatively homogenous cohort of young individuals without significant comorbidities and provides a comprehensive understanding of dynamic and nuanced temporal changes in ADAR editing patterns, across three distinct stages of SARS-CoV-2 infection. While this cohort simplifies analysis by reducing potential confounding factors, such as age-related variations and comorbidities, it is important to acknowledge its limitations. First, ADAR mediated editing shows significant variability across different tissues often associated with differences in the expression of transcripts incorporating the edit (Huntley et al., 2016). Moreover, the sequencing data used in our study were derived from whole blood samples, and thus, they may not capture the tissue-specific molecular changes such as those occurring in the respiratory tract (primary site of infection) or organs directly or indirectly affected by infection such as the brain (as observed in infections with mycobacterium tuberculosis) (Brighenti & Andersson, 2012). Next, sex-specific differences are widely reported during SARS-CoV-2 infection, including within the original cohort of the CHARM dataset, as well as in ADAR editing patterns are increasingly being recognized in various disease states (Sauerwald et al., 2022). Therefore, the possibility of sex as a variable influencing ADAR editing patterns cannot be ruled out. Our dataset also suffers from technical limitations, namely the number and levels of identified editing site correlates with the sequencing depth of the input samples. As a sequencing depth of around 80-100 million paired-end reads is recommended for Illumina sequencing (Diroma et al., 2019), our samples had an average sequencing depth of 25 million reads (Sauerwald et al., 2022). This low sequencing depth might have resulted in an underestimation of editing changes in our dataset. Therefore, future studies with higher sequencing depth are warranted to understand the full spectrum and range of changes in editing during SARS-CoV-2 infection.

## Conclusions

Overall, our findings show the dynamic nature of SARS-CoV-2 induced changes in ADAR1 expression and activity across distinct stages of infection. By using samples from same individuals across three stages of infection, along with our stringent filtering criteria, we identify unique ADAR editing sites across all three stages, resulting from either loss or introduction of new sites. Noteworthy, our findings suggest a consistent pattern of ADAR expression and editing among individuals during active SARS-CoV-2 infection, where ADAR1 expression is elevated and likely translates to downstream alterations in ADAR editing patterns. However, post infection, while ADAR1 expression consistently returns to pre-infection levels, ADAR editing patterns remain dysregulated in a subset of individuals, attesting to the heterogenous molecular patterns widely observed in individuals post-SARS-CoV-2 infection. While our study suggests persistent dysregulation of ADAR editing patterns in a subset of individuals post-viral clearance, the mechanistic significance and implications of this dysregulation, including in SARS-CoV-2 infection sequelae needs to be elucidated.

## Methods

### RNA-sequencing dataset from mild COVID-19 patients across three distinct stages of SARS-CoV-2 infection

The dataset used in this study includes whole-blood RNA sequencing samples from a subset of individuals from the COVID-19 Health Action Response for Marines (CHARM) study, which are publicly available at NCBI-GEO under the accession GSE198449 (BioProject PRJNA815324) (Sauerwald et al., 2022). Briefly, CHARM was a prospective study that tracked the progression of SARS-CoV-2 infection among marine recruits entering military training from May 11,2020 to November 2, 2020. The cohort consisted of young, age-matched (average age 19.1 ± 1.9 years), physically fit individuals without documented comorbidities. Individuals in the cohort mostly developed mildly symptomatic disease with symptoms including abdominal pain, chills, cough, decrease in smell and taste, diarrhea, fatigue, fever, headache, muscle ache, nausea, runny nose, sore throat, and shortness of breath. Participants were tested for the presence of SARS-CoV-2 infection across different stages of SARS-CoV-2 infection using serial nasal swab PCR. Results of PCR testing were used to annotate RNA-sequencing samples (read length 100 bp) to distinct stages of infection. The study deposited 1858 RNA-sequencing samples from 475 individuals.

We selected a subset of 45 individuals from the original cohort for whom RNA-sequencing samples were available across three distinct stages of SARS-CoV-2 infection; pre-, mid-, and post-infection (Supplementary Table 9). All the samples labelled “control” at day T0, representing individuals tested negative for SARS-CoV-2 on PCR, were categorized as pre-infection samples. Subsequently, all the samples labelled “mid”, representing samples tested PCR positive for SARS-CoV-2, on day T35 in the original dataset were selected as mid-infection samples. Selection of Day T35 for mid-infection samples was done to account for the incubation period of SARS-CoV-2 which has a mean incubation period of 6.57 days ranging from 1.80 to 18.87 days (Wu et al., 2022). Additionally, selection of day T35 for mid-infection samples allowed us to incorporate maximum number of individuals in our study. Finally, all the samples labelled “post”, representing samples that returned to negative status for presence of SARS-CoV-2 on PCR, in the original dataset, with an average gap of 80.5 days (T56 -T105), were selected as post-infection samples.

### RNA seq data analysis, variant calling and identification of ADAR editing sites

The computational pipeline Automated Isoform Diversity Detector (AIDD) (Plonski et al., 2020) was used to map, assemble, and perform variant calling on RNA seq datasets to identify ADAR editing sites. Briefly, RNA sequencing samples were downloaded from NCBI sequence read archive (NCBI-SRA), after performing quality control of raw reads using FASTQC (https://www.bioinformatics.babraham.ac.uk/projects/fastqc/), HISAT2 (Kim et al., 2015) was used for alignment of reads to the reference genome GRCh37 (human reference Ensembl built release 75). An average of ∼ 53.59 million paired reads per sample for the 135 samples from 45 patients were aligned to the genome (Table 2). Supplementary Table 10 details the number mapped reads for each sample. Subsequently, Stringtie (Pertea et al., 2015) was used to perform transcriptome assembly. Expression of genes, including ADARs, were estimated in Transcripts Per Kilobase Million (TPM). Stringtie generated ballgown files contain gene and transcript level information, a custom python script was applied to the ballgown files to generate gene and transcript level raw count files. DESeq2 (Love et al., 2014) was used to perform differential gene expression analysis using raw gene- and transcript-counts. p adjusted was calculated using the Benjamini-Hochberg (BH) procedure to control for False Discovery Rate (FDR). Expression of ADAR isoforms were analyzed using DESeq2 normalized raw transcript counts. Differentially expressed genes were annotated using the R package biomaRT (Durinck et al., 2009). Further the aligned and annotated BAM files were used for Variant calling using GATK haplotype caller (McKenna et al., 2010). The process followed the recommended GATK best practices, as outlined in Plonski et al. (2020), to infer putative RNA editing events (Plonski et al., 2020). Generated VCF files were used to determine the total number of putative ADAR editing sites, which includes both A-to-G and T(U)-to-C edits, after filtering for known SNPs from NCBI database of single nucleotide polymorphisms and potential polymorphic sites (Sherry et al., 2001)(sites with multiple nucleotide outcomes). Counts from the bam-read count (Khanna et al., 2021) were used to determine variant frequency counts to calculate per site editing levels. Total number of putative ADAR edits per sample were normalized for sequencing depth.

### Identification of consistent and unique ADAR editing sites

We identified ADAR editing sites consistently edited in pre-, mid-, and post-infection samples as well editing sites uniquely edited across all three stages of infection (Supplementary table 6). Accounting for inter-individuals variability in ADAR editing (O’Neil et al., 2017), consistently edited sites were defined as those A-to-G and T-to-C substitutions that satisfied the following criteria; had at least 100 reads aligned (stack depth) at each ADAR edited site (A-to-G or T-to-C substitutions), were present in at least 20% of samples for a given stage of infection, with an editing level (defined as the number of G reads for a reference of A and the number of C reads, reference site on complimentary strand, for a reference of T to the total number of reads aligned to that site) of at least 2% or greater, and had a GATK mapping quality of at least 40. Next, we compared consistently edited sites between pre- and mid-infection samples and between pre- and post-post-infection samples to identify ADAR editing sites unique to a stage of infection. The identified unique sites were annotated using REDIportal database V2.0 (Picardi et al., 2017), a comprehensive repository of more than 4.5 million ADAR editing events from humans. Reactome pathway analysis (Fabregat et al., 2017) was used to explore pathways enriched among sites uniquely edited pre-, mid-, and post-infection.

### Global ADAR editing levels

To analyze global changes in the editing levels, we calculated overall ADAR editing levels per sample using methods previously described (Breen et al., 2019; Tan et al., 2017). Briefly, AIDD generated VCF files containing all possible base substitutions were filtered to include only ADAR editing sites (A-to-G and T-to-C), with number of aligned reads greater than or equal to 100. To filter out potential false positives, the remaining sites were further filtered to include sites previously defined in REDIportal (Picardi et al., 2017). This was done to include only high confidence previously identified ADAR editing sites. Next, overall ADAR editing levels were calculated as the ratio of total number of edited reads aligned to ADAR editing sites present in the sample to the total number of all reads aligned to ADAR editing sites in that sample. Finally, we performed unsupervised *K-*means clustering to analyze whether dysregulation in ADAR editing patterns follows distinct patterns among individuals both mid- and post-infection. We performed two separate clustering analyses on overall ADAR editing levels of samples from pre- to mid-infection and from pre- to post-infection. The optimal number of clusters was determined by using the elbow method, and a value of K =2 was selected, representing the lowest within-group sum of squares among the clusters. The analysis was performed using ‘kmeans’ function from base R.

### Statistical analysis and visualization

All the comparisons were made keeping pre-infection as the baseline level and paired t-test was used to identify changes in expression of all three ADARs, ADAR isoforms, total number of ADAR editing sites and overall ADAR editing levels between distinct stages of infection using the “stat_compare_means” function from the ggpubr (version 0.6.0 - https://rpkgs.datanovia.com/ggpubr/) package in R. Code for upset plot (Figure 3B) has been adopted from DOI: 10.5281/zenodo.7555525. Supplementary materials and code are shared at https://github.com/RNAdetective/Temporal_editing_in_SARSCoV2.

## List of abbreviations

ADAR: Adenosine Deaminases Acting on RNA
IFN: Interferon
ISG: Interferon stimulated gene
ISRE: Interferon-stimulated response element
UTR: Untranslated region
SARS-CoV-2: Severe acute respiratory syndrome coronavirus 2
COVID-19: Coronavirus disease 2019
ssRNA: single stranded RNA
dsRNA: double stranded RNA
TPM: Transcripts per million
CHARM: COVID-19 Health Action Response for Marines

## Supplementary Materials

## List of supplementary tables and figures

Supp_Table_1. Supplemental_Table_1A: DESeq2 differential gene expression analysis results comparing Pre-infection samples to Mid-infection samples, filtered for log2Fold Change > |0.58| (Fold Change > 1.5) and an adjusted P value < 0.05.

Supplemental_Table_1B: DESeq2 differential gene expression analysis results comparing Pre-infection samples to Post-infection samples, filtered for log2Fold Change > |0.58| (Fold Change > 1.5) and an adjusted P value < 0.05.

Supplementary Table_1C: List of interferon stimulated genes differentially expressed mid-SARS-CoV-2 infection compared to pre-infection.

Supplemental_Table_1D: DESeq2 differential gene expression analysis results comparing Pre-infection samples to Mid-infection samples, filtered for log2Fold Change > |1| (Fold Change > 2) and an adjusted P value < 0.05.

Supplemental_Table_1E: DESeq2 differential gene expression analysis results comparing Pre-infection samples to Post-infection samples, filtered for log2Fold Change > |1| (Fold Change > 2) and an adjusted P value < 0.05.

Supp_Table_2. Supplemental_Table_2A: Reactome pathway over representation (OR) analysis results for genes differentially expressed mid-infection compared to pre-infection.

Supplemental_Table_2B: Reactome pathway over representation (OR) analysis results for genes differentially expressed post-infection compared to pre-infection.

Supplemental_Table_2C: Reactome pathway over representation (OR) analysis results for genes differentially expressed mid-infection compared to pre-infection (using stricter cut off Log2FC > |1| & adj p value <0.05.

Supp_Table_3. Supplemental_Table_3A: ADAR1 expression in TPM for each patient across pre-,mid-, and post-infection.

Supplemental_Table_3B: ADAR2/ADARb1 expression in TPM for each patient across pre-,mid-, and post-infection.

Supplemental_Table_3C: ADAR3/ADARb2 expression in TPM for each patient across pre-,mid-, and post-infection.

Supp_Table_4. Supplemental_Table_4A: Transcript level ADAR expression as DESeq2 normalized counts. Isoform transcript Ids were retrieved from Ensembl.

Supplemental_Table_4B: Transcript level ADAR expression data from DESeq2.

Supp_Table_5. Supplemental_Table_5: Total number of putative ADAR editing sites for each patient across pre-, mid-, and post-infection stages of SARS-CoV-2 infection.

Supp_Table_6. Supplemental_Table_6A: List of sites consistently edited pre-, mid-, and post-infection.

Supplemental_Table_6B: List of sites uniquely edited pre-, mid-, and post-infection. Supplemental_Table_6C: REDIportal annotated pre-infection unique sites.

Supplemental_Table_6D: REDIportal annotated mid-infection unique sites. Supplemental_Table_6E: REDIportal annotated post-infection unique sites.

Supp_Table_7. Supplemental_Table_7: Reactome pathways over representation (OR) analysis of genes incorporating confirmed unique editing sites (A) pre-, (B) mid-, and (C) post-infection.

Supp_Table_8. Supplemental_Table _8: Patient-wise overall ADAR editing levels across pre-, mid-, and post-infection stages of SARS-CoV-2 infection.

Supp_Table_9. Supplemental_Table_9: List of 45 individuals selected from the initial cohort of CHARM dataset with samples available across pre-, mid-, and post-infection stages.

Supp_Table_10. Supplemental_Table_10: Metrics of analyzed RNA-seq data from Bioproject PRJNA815324; GSE198449 (Sauerwald et al.,2022).

**Supplemental Figure 1:**
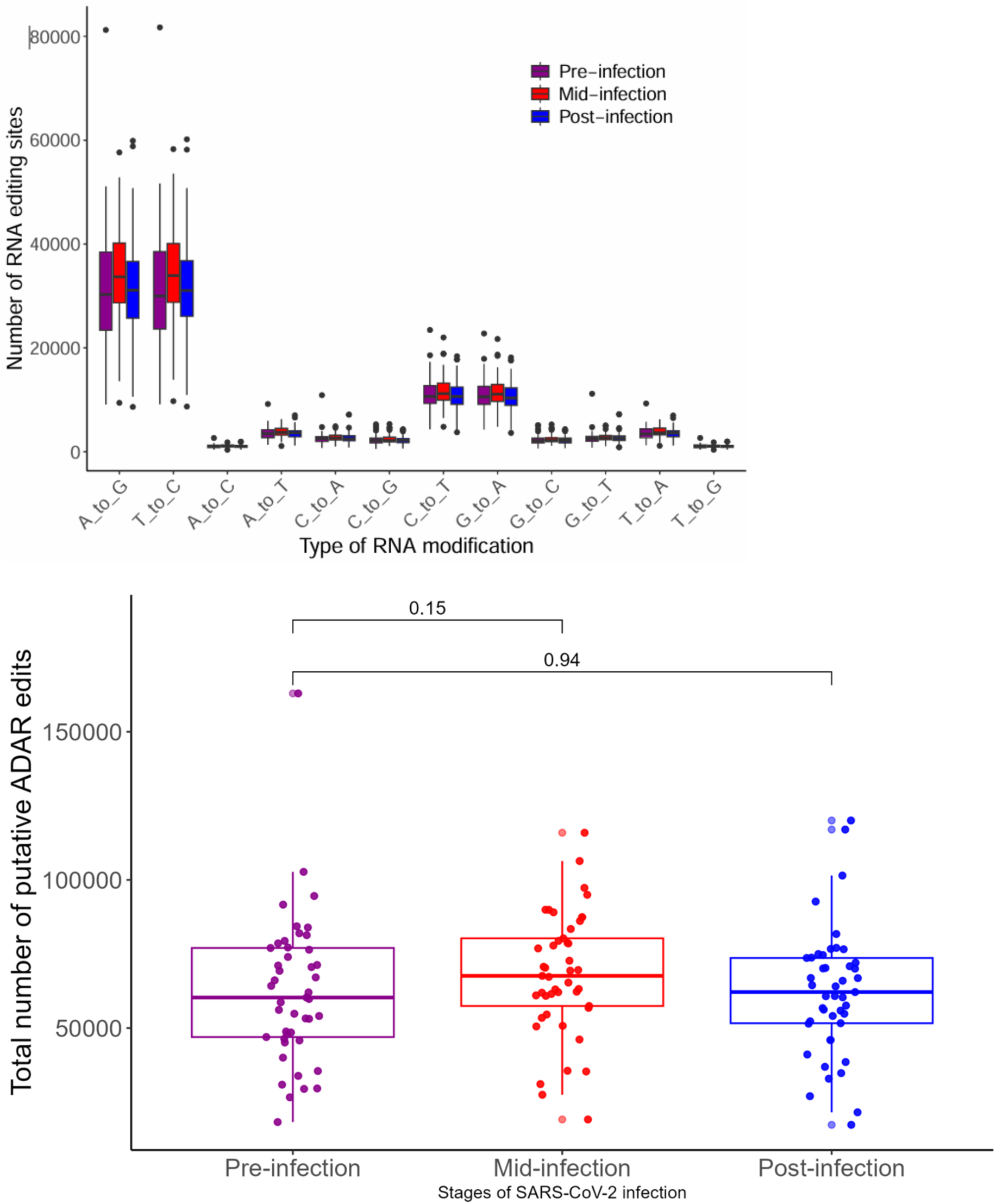
Alterations in total number of putative ADAR edits mid- and post-infection compared to pre-infection. A-to-G and T-to-C substitution representing ADAR potential ADAR edits were the most abundant form of substitutions across all three stages of infection (A). While the total number of putative ADAR edits that includes both A-to-G and T-to-C substitutions increased mid-infection, the change did not achieve statistical significance (paired t-test, p value = 0.15). Similarly, there was no significant change in the number of putative ADAR edits post-infection compared to pre-infection (paired t-test, p value = 0.94) (B).

**Supplemental Figure 2:**
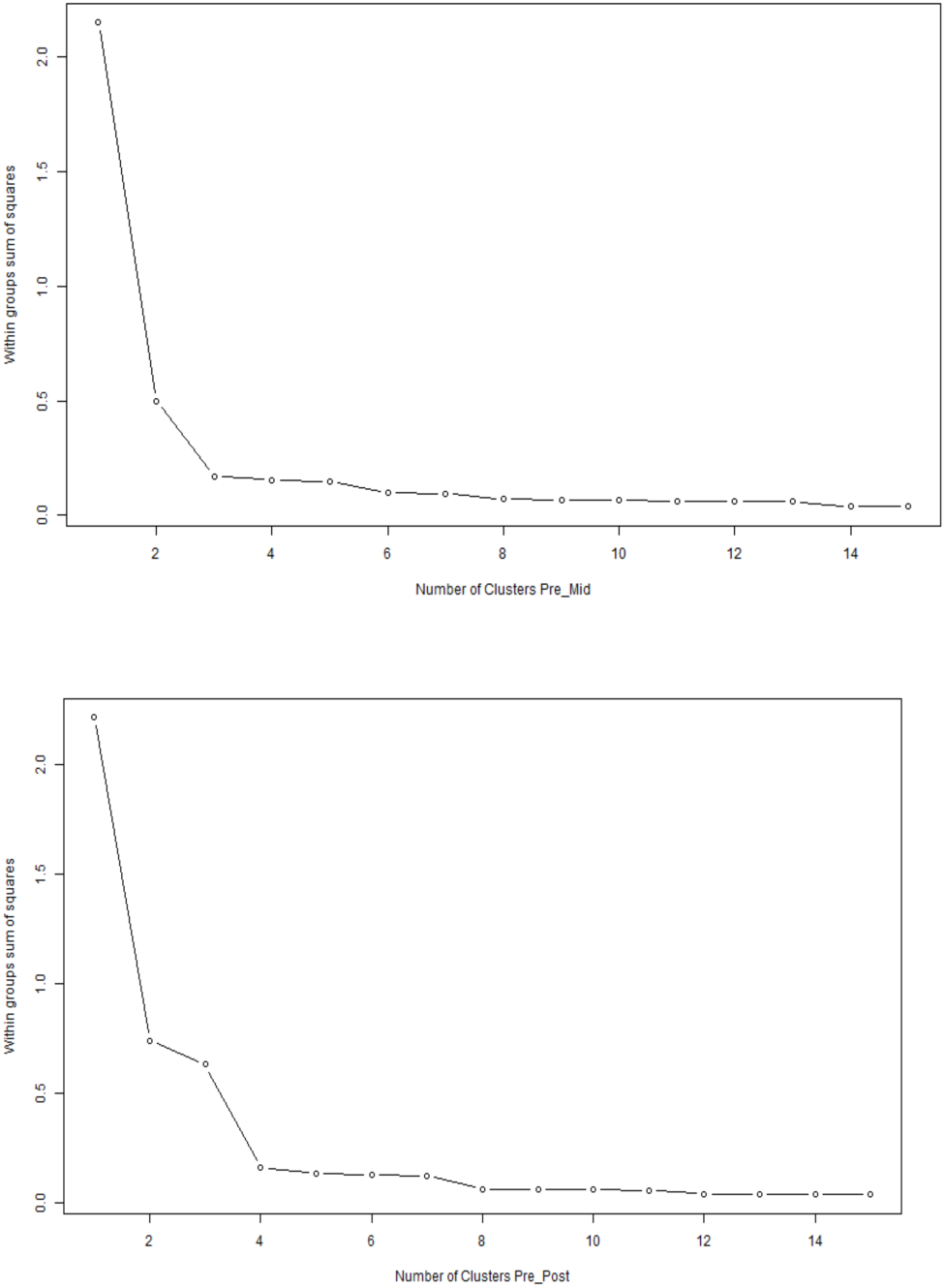
Elbow plots showing the total number of potential groups based on overall ADAR editing values in individuals’ pre- and mid-infection (2A) and pre- and post-infection (2B).

